# Dynamic ecological interactions of two quarantine-concern *Ralstonia* strains in river water and plants

**DOI:** 10.1101/2025.10.21.683785

**Authors:** Bridget S. O’Banion, Jake A. Criscuolo, Hanlei Li, Addison Alderdice, Caitilyn Allen

**Affiliations:** Department of Plant Pathology, University of Wisconsin-Madison, Madison, USA

**Keywords:** competitive fitness, aquatic microbial ecology, *Ralstonia solanacearum*, *Ralstonia pseudosolanacearum*, bacterial wilt, potato brown rot, U.S. Select Agent pathogen

## Abstract

Plant pathogenic *Ralstonia* belonging to the IIB-1 (“race 3 biovar 2”) and I-33 (rose) subgroups are emerging quarantine and biosecurity threats. Both strains have been introduced to Europe, where they persist in weedy plants and surface water and cause occasional costly disease outbreaks. We combined *in planta, in vitro*, and environmental water microcosm experiments to determine if these two concerning strains are likely to co-exist in environments where they have become established or if one might be expected to displace the other. Using a representative strain from each subgroup we investigated the dynamics and fitness of these two *Ralstonia* pathogens across ecologically relevant environments. Interactions between the strains were context dependent: the presence of a competing strain had little impact on bacterial survival in river water microcosms, but the I-33 strain had a fitness advantage in wilt susceptible tomato plants. We found no evidence of direct growth inhibition by either strain *in vitro*. The IIB-1 strain persisted longer than I-33 in cool temperature river water microcosms. Warmer temperatures extended the culturability of both strains, which may be important as climate change warms surface water globally. Additionally, *Ralstonia* strains persisting in 20°C water microcosms for 6 months were still able to cause disease in tomato plants. Together, our results provide useful insight into the dynamics of these two strains in environments where they are currently established, which may inform management practices moving forward.

## Introduction

Bacterial wilt caused by plant pathogenic *Ralstonia* is a destructive crop disease that begins with colonization of plant vascular systems through the roots, leading to rapid wilting and death. The *R. solanacearum* species complex (RSSC) consists of three species: *R. solanacearum* (*Rs*, phylotype II), *R. pseudosolanacearum* (Rps, phylotype I and III), and *R. syzygii* (*Rsy*, phylotype IV) (Safni *et al*. 2014; Prior *et al*. 2016). The RSSC has an unusually broad host range, infecting species in over 250 diverse plant families, and bacterial wilt is a limiting factor for major crops like potato, tomato, banana, tobacco, and peanut (Vailleau and Genin 2023). These gram-negative soil-dwelling bacteria persist in plants and in environmental reservoirs like soil and water. They can asymptomatically colonize plants to survive harsh environmental conditions. These reservoirs have made it difficult to eradicate *Ralstonia* introductions or contain bacterial wilt outbreaks.

A near-clonal *Rs* lineage known as IIB-1 (phylotype II, sequevar 1) causes brown rot disease of potato, especially in temperate zones and the cool highland tropics (Allen, C., A. Kelman, and E.R. French 2001). Also known as Race 3 biovar 2, the IIB-1 lineage co-evolved with the potato in the Andean highlands and has been globally disseminated in latently infected seed potato tubers (Clarke *et al*. 2015; Dewberry *et al*. 2024). In addition to causing major yield losses on potato, a key carbohydrate crop, the IIB-1 lineage has also inflicted economic damages associated with quarantine enforcement and eradication efforts. It is a highly regulated Select Agent in the U.S. and a quarantine pathogen in Europe, where it was accidentally introduced in the 1990s and has persisted in soil, surface water, and weeds (Janse *et al*. 2004; Parkinson *et al*. 2013; Álvarez, López and Biosca 2022). Eradication efforts continue in Europe, and testing and certification of seed potato tubers help counteract *Ralstonia* spread (European Food Safety Authority (EFSA) *et al*. 2025). Nonetheless, IIB-1 is unfortunately now widely distributed across the European landscape.

*Rps* phylotype I, long considered a tropical pathogen, was first reported in Europe in 2015. An *Rps* strain belonging to phylotype I sequevar 33 strain (I-33) was isolated from ornamental rose in a Dutch greenhouse facility (Tjou-Tam-Sin *et al*. 2017a). Sequence analysis indicates that this strain, which has proven difficult to eradicate, has infected rose in several European and Asian countries (EPPO 2016, 2017). Concerningly, *Rps* I-33 was recently detected in geographically distinct surface waterways in the Netherlands and Hungary (EPPO 2021a, 2022a; Vogelaar *et al*. 2023). Some surveys detected *Rps* phylotype I together with *Rs* IIB-1. Further, additional genetically distinct *Rps* phylotype I strains have been detected in field- and greenhouse-grown ginger, turmeric, cucumber, tomatoes, and potatoes in multiple European countries (EPPO 2021b, 2022b, 2022c, 2023; Holeva *et al*. 2024).

It is now clear that multiple diverse strains of plant pathogenic *Ralstonia* are present in Europe. These bacteria are likely to interact in surface waterways, irrigation systems, and in wild hosts like the common semi-aquatic weed bittersweet nightshade (*Solanum dulcamara*). However, it is not known how or if they affect each other. We previously showed that *Rps* I-18 strain GMI1000 outcompetes IIB-1 strain UW551 in plants at tropical temperatures, but at cooler temperatures the IIB-1 strain had much higher competitive fitness, nearly excluding *Rps* strains from co-inoculated tomato plants (Huerta, Milling and Allen 2015). To better predict how the *Rs* and *Rps* strains now established in Europe interact in the landscape, we used a representative strain from each of these near-clonal groups for a combination of *in planta, in vitro*, and environmental water microcosm experiments. The two strains interacted minimally in water, but I-33 had a fitness advantage in wilt susceptible tomato plants. Warmer water temperatures extended the culturability of both strains, and the IIB-1 strain persisted for longer in cool temperature river water microcosms. Interestingly, the temperature of a 6 month aquatic incubation determined whether the *Ralstonia* cells remained virulent in tomato plants.

## Materials and Methods

### General bacterial strains/maintenance

*R. solanacearum* strain UW551-GFP-tet^R^ is a typical IIB-1 (“Race 3 biovar 2”) brown rot pandemic strain. It encodes a constitutively expressed green fluorescent protein (GFP) fluor and tetracycline resistance but retains wild-type virulence (Swanson *et al*. 2005; Hayes, MacIntyre and Allen 2017; Criscuolo *et al*. 2025). *R. pseudosolanacearum* strain UW770-rif^R^ is a I-33 strain originally isolated from rose in the Netherlands in 2015 and is also known as IPO4001 and PD7123 (Tjou-Tam-Sin *et al*. 2017b; van der Wolf *et al*. 2022). The strain carries spontaneous rifampicin resistance and retains wild-type virulence (van der Wolf *et al*. 2022). *Ralstonia* strains were cultured and stored as described (Khokhani *et al*. 2018).

### River water collection and preparation

River water was collected from the Yahara River in Madison, Wisconsin (43°09’09.0”N 89°24’02.9”W) in August 2024 near USGS monitoring site ID 05427850. Water for each of the three microcosm replicates was independently collected at least 1 week apart. Water was collected in the evening and stored at 4°C in 1 liter plastic Nalgene bottles until sterilization the following day. Each batch of water was filter-sterilized with a 0.22 μm filter to remove debris and microbial cells, autoclaved for 20 minutes at 121°C to destroy viruses, and filter-sterilized once more to remove precipitate. The water pH was 8.06, 8.09, and 8.14 for replicates 1, 2, and 3, respectively. The sterilized water was stored at 4°C until the next day, when it was used for experiments. Historical climate data was accessed via the National Oceanic and Atmospheric Administration’s Climate Data Online Search (https://www.ncdc.noaa.gov/cdo-web/search). The daily precipitation and temperature summaries from August 4th to 26th, 2024 for surrounding and upstream zip codes used for analyses.

### Water microcosm set-up

A single colony of each strain was picked from a rich casamino acids-peptone-glucose (CPG) medium agar plate and used to inoculate 100 mL of CPG broth. Antibiotics were added to CPG as needed: 25 mg/L tetracycline for UW551-GFP-tet and 25 mg/L rifampicin for UW770. The next day, the O.D._600_ of cultures was monitored until each strain reached early log phase growth (OD_600_ 0.1-0.2). One hundred mL of each cell suspension was then centrifuged at 2,500 x *g* for 5 minutes. The supernatant was removed and bacterial cells were resuspended in sterile water before repeating this washing process once more. The washed bacterial suspensions were then used to inoculate the sterile river water to a final O.D._600_ of 0.01 (corresponding to ∼1×10^7^ CFU/mL) using either UW551-GFP-tet^R^, UW770-rif^R^, or a 1:1 mixture of each. A 100 μL sub-sample of all inoculated river water treatments was immediately taken and used to quantify the initial bacterial population. Twenty-five mL of the inoculated river water was then aliquoted into each of twelve 50 mL falcon tubes for each treatment. All tubes were incubated at room temperature for 48 h to allow the cells to consume labile nutrients. A sub-sample was collected from each tube after this initial bloom to quantify cells prior to long-term storage at experimental temperature conditions. All tubes for a respective temperature were placed in a conical tube rack nested inside a black tray, covered with a secondary black tray, and statically incubated at either 4°, 12°, or 20°C.

### Water microcosm sampling

Each week, tubes were briefly removed from their respective temperature storage conditions for sampling. Tubes were mixed with 2 to 3 vigorously inversions and 100 μL was removed for bacterial quantification. These sub-samples were serially diluted in water and plated on solid CPG containing the appropriate antibiotics. Sub-samples from co-inoculated microcosms were plated on both CPG + rifampicin and CPG + tetracycline to enumerate both strains. Plates were incubated at 28°C for 48-72 h until countable colonies formed. Beginning at week 3 of sampling for each replicate, a suspension of commercial catalase was spread on plates immediately before plating samples to reduce oxidative stress and help facilitate cellular growth. During late stages of the experiment, sampling frequency was reduced due to cell numbers remaining stable. Once CFUs were no longer detectable within a specific set of tubes, sampling waas discontinued.

### Plant maintenance and growth conditions

Tomato plants (wilt-susceptible cv. ‘Bonny Best’) were grown in a climate chamber at 28°C with a 12-hour photoperiod as described (Khokhani *et al*. 2018). Briefly, seeds were planted in a growing mix and transplanted to 4-inch pots at 14 days old. Once transplanted, they were watered with a rotation of water and 1/2-strength Hoagland’s solution.

### Virulence and stem colonization

Plant assays were performed as described (Khokhani *et al*. 2018). Briefly, 21 to 22 day-old plants were petiole-inoculated with 2 µL of a 10^6^ CFU/mL suspension of UW551-GFP-tet^R^, UW770-rif^R^, or a 1:1 mixture of each. For virulence assays, symptom development was rated for 14 days on a 0-4 disease index scale. To quantify stem colonization, approximately 100 mg of stem directly below the cut petiole was sampled three days post-infection and homogenized in water. Samples were diluted and plated on solid CPG + appropriate antibiotics to quantify CFU/gram. To qualitatively determine colonization (stamp assays), plants were de-topped directly above the site of inoculation at the end of the monitoring period using a razor blade. The blunt end of the de-topped foliage was then pressed firmly against a solid CPG agar plate. The plates were incubated for up to a week to monitor growth of *Ralstonia* cells, which were identified by their distinctive fluidal colony morphology.

For plants inoculated with water-adapted *Ralstonia*, 1 mL was removed from the aquatic microcosms (approximately 25 weeks post-inoculation) and centrifuged at 8,000 x g for 10 minutes. The water was removed and the pellet was resuspended in 10 uL of water. This was then used to inoculate 3 individual plants. Each of the three aquatic microcosm replicates was used to inoculate 3 separate plants (n=9 per treatment).

### Cell-free supernatant assays

UW770 and UW551 cultures were grown overnight in CPG broth. The following day, 1 mL of the turbid culture was centrifuged at 8,000 x g for 5 minutes to pellet the cells. The supernatant was then sterilized by passing it through a 0.22 um filter. Separately, molten semi-solid CPG agar (cooled to ∼45°C) was mixed with washed *Ralstonia* cells to a density of OD_600_ 0.01 and poured into petri plates. This facilitates the growth of a solid lawn of cells. After drying, 5 uL of the filter-sterilized supernatant was spotted on top of the agar and allowed to dry. Plates were made with UW770 as the lawn and UW551 as the lawn. For each strain as a lawn, the supernatant spots included the following: supernatant produced by self, supernatant produced by the other strain, sterile CPG media, and water. Duplicate spots for each treatment and control were placed on each plate. The experiment was repeated 3 separate times.

## Results

### Temperature has a strain-dependent impact on RSSC fitness in river water

Freshwater serves as a critical reservoir for *Ralstonia* globally. We therefore tested the persistence (measured as culturable cells) of these two strains over a 6 month incubation in sterilized river water microcosms incubated at 4**°**C, 12**°**C, and 20**°**C, which are likely water temperatures in various surface waterways in Europe over the seasons. We hypothesized IIB-1 strain UW551 would survive better than I-33 strain UW770 at all of these temperatures due to its evolutionary history in cooler regions. At both 4**°**C and 12**°**C, UW551 indeed persisted at higher levels than UW770 (Figure 1A-B). Strain UW770 became undetectable via plating 4 weeks earlier than UW551 in the 4**°**C microcosms and 10 weeks earlier in the 12**°**C microcosms. Both strains eventually become non-culturable at these temperatures within the time-frame of our experiment. Neither strain could be detected via plating within 2 months at 4**°**C and 5 months at 12**°**C (Figure 1A-B). Overall, both *Ralstonia* persisted longer at 20**°**C, with both strains still detectable at >10^5^ CFU/mL after 6 months with indistinguishable population sizes (Figure 1C).

**Figure 1.**
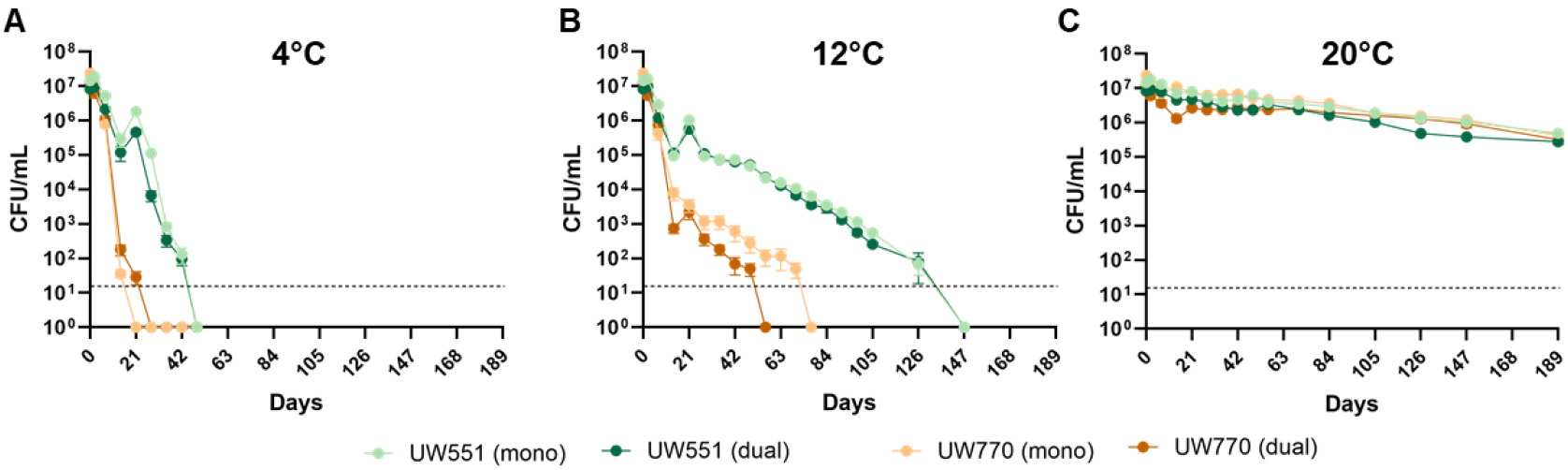
*Ralstonia* strain culturability across aquatic microcosm treatments. *R. solanacearum* **IIB-1 strain** UW551 (gold) and *R. pseudosolanacearum* I-33 strain UW770 (green) were inoculated singly (mono) or dually (dual) into river water microcosms and incubated at either A) 4°C, B) 12°C, or C) 20°C for 6 months. Lines represent the average of 3 individual tubes for 3 biological replicates (n=9). Bars represent the SEM at each timepoint. The “Dual” lines represent the CFUs of each respective strain from the dual-inoculated microcosms. A single sub-sample from the dual-inoculated tubes was dilution plated on CPG + appropriate antibiotic to enumerate each strain individually. Plates used for sampling after 21 days also contained catalase. The panels represent an increasing temperature gradient from left to right.

### UW551 and UW770 exhibit minimal interaction in water microcosms

In addition to measuring culturability over time when inoculated singly into water microcosms, we also co-inoculated UW551 and UW770 together. Members of the RSSC are known to antagonize each other with highly specific bacteriocins. We therefore hypothesized that co-inoculations would have deleterious effects on the persistence of the strains, as compared to their respective mono-inoculated microcosms. Instead, we found that the presence of a potential competitor did not dramatically impact the persistence of either strain (Figure 1A-C). UW551 and UW770 became undetectable via plating at the same, or similar, timepoints in the 4**°**C and 12**°**C microcosms regardless of whether they were inoculated alone or together. There is a small separation between the mono- and dual-UW770 trajectories at 12**°**C. However, this difference is driven by a single replicate (Figure S1, blue line (replicate 3)).

We next analyzed publicly-available climate data for the days surrounding our river sampling timepoints. The days leading up to water sampling for replicate 3 had the longest period of drought, and the daily temperature was highest on the day of collection (Figure 2). Climatic variations can lead to changes in the physiochemical properties of water. We therefore hypothesized that the concentration of specific ions or nutrients might explain this variation. To test this, we measured a suite of anions, cations, and total organic and inorganic carbon in each of the three water samples. However, none of the measured variables showed a notable difference specifically for replicate 3 (Table S1).

**Figure 2.**
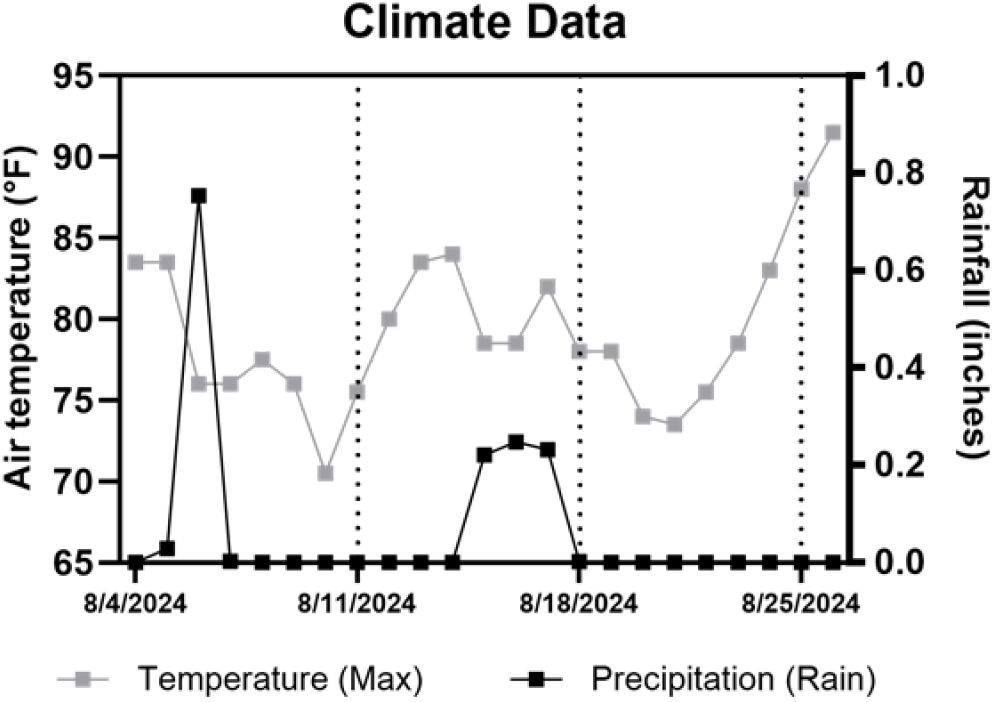
Climate data for regional watershed during sampling season. The maximum air temperature (gray) and rainfall (black) records from surrounding USGS monitoring locations were averaged together for days surrounding the sampling season. Squares represent the mean value for each day. Horizontal dotted lines represent the three days of water collection for the aquatic microcosms. Replicate 1 was collected on August 11th, replicate 2 was collected on August 18th, and replicate 3 was collected on August 25th.

### UW770 has a significant fitness advantage over UW551 in tomato stems at 28°C

We were next curious how these two strains interact with each other in a common plant host. To investigate this, tomato plants were petiole-inoculated with UW551, UW770, or a 1:1 mixture of both strains and kept at 28°C. Stem sections were collected three days post-inoculation to quantify colonization. When the strains were inoculated individually, UW770 colonized stems at significantly higher densities than UW551 (Figure 3A). In the dual-inoculations, UW770 colonized at significantly higher levels than UW551 (Figure 3B-C). Of the 36 dual-inoculated plants, 94.5% were dominated by UW770, while UW551 dominated the stem in only 5.5% of the plants (n=2). On average, a single infected plant stem contained 99.9% more UW770 cells than UW551. We tested the hypothesis that this competition was mediated by the production of antimicrobial compounds, such as bacteriocins. However, an *in vitro* supernatant assay provided no evidence that secreted compounds produced by either strain had a deleterious effect on the other strain’s growth (data not shown).

**Figure 3.**
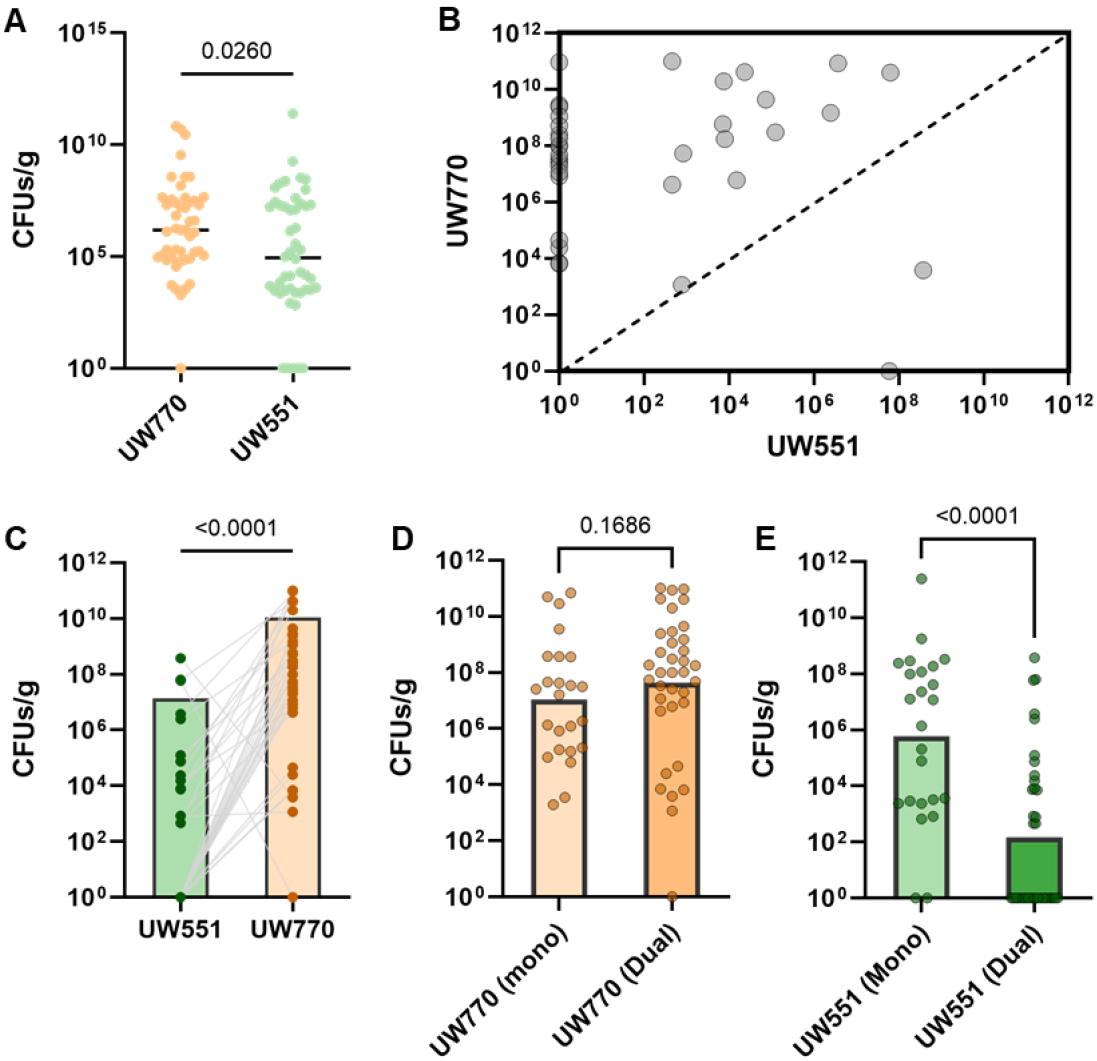
*Ralstonia* phylotype I-33 strain UW770 has a fitness advantage in wilt-susceptible tomato plants at 28°C. A) Stem colonization of each strain when inoculated individually into plants via petiole-inoculation. Stems were harvested 3 days post-inoculation. P-value determined by Mann-Whitney test. B-C) Stem colonization of each strain when co-inoculated into plants at a 1:1 ratio via petiole inoculation. Stems were harvested 3 days post-inoculation and homogenate was plated on CPG + antibiotics to separately measure the colonization of each strain. In panel B, the dotted line represents an equivalent population of each strain. Dots that fall in the upper left quadrant represent individual plants with a higher level of UW770, while dots in the lower right quadrant represent individual plants with a higher level of UW551. In panel C, the population level of each strain within a single individual plant is represented by two dots connected by a line. The green dots represent the population of UW551 cells and the orange dots represent the population of UW770 cells. A green dot connected by a gray line to an orange dot indicates the respective UW551 and UW770 populations within a single stem. The data in panel C is the same data shown in panel B. The P-value in panel C was determined by a Wilcoxon matched-pairs signed rank test. Stem colonization of UW 770 (D) and UW551 (E) in mono vs. co-inoculation. Each dot represents a single plant. P-values determined by Mann-Whitney test.

The mono-colonization level of UW770 was similar to its colonization outcomes in dual-inoculated plants (Figure 3D). On the other hand, the colonization level of UW551 in a competitive environment was significantly less than its colonization in a non-competitive environment (Figure 3E). In many dual-inoculated plants, UW551 cells were not detectable via plating. Intrigued by the *in planta* dominance of UW770 following a 1:1 inoculation mixture (Figure S2A), we increased the inoculum ratio of UW551. We observed a similar dominance of UW770 when UW551 cells made up 70% of the initial inoculum (Figure S2B). However, when we increased UW551’s ratio to 98% of the initial inoculum, we observed UW551 dominance in the harvested stems (Figure S2C).

### Water-adaptation modestly influences the stem colonization of UW770 and UW551

We were next curious if 6 months in river water changed the colonization or virulence potential of each strain, as compared to the same strain kept in typical laboratory storage conditions (frozen in glycerol). We therefore used concentrated water from the 20°C water microcosms or typical washed and diluted overnight cultures to perform colonization and virulence assays. To test colonization, we petiole-inoculated susceptible tomato plants and harvested stem sections three days later. We found that water-adaptation did not change the stem colonization levels of UW551 as compared to cells stored in typical laboratory storage conditions (Figure 4A). However, water-adaptation caused a significant decrease in stem colonization levels of UW770 (Figure 4A).

**Figure 4.**
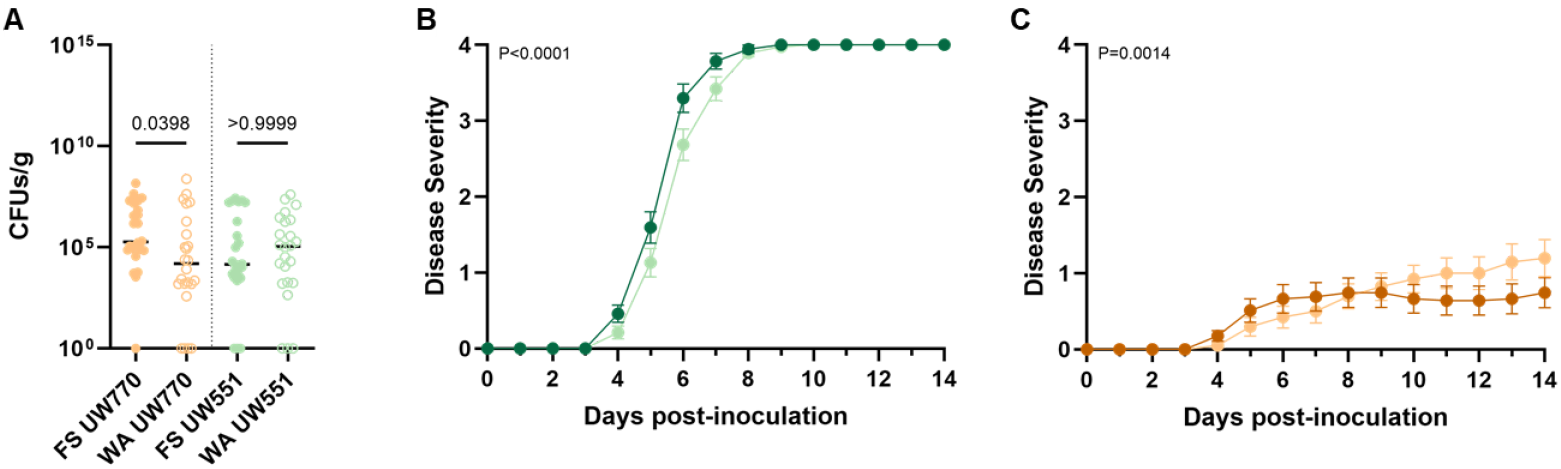
Long-term persistence in 20°C water microcosms has modest, strain-specific effects on plant colonization and virulence potential of cells. A) Stem colonization of each strain when inoculated individually into plants via petiole-inoculation. FS = freezer stock, WA = water-adjusted. Stems were harvested 3 days post-inoculation. Each dot represents a single plant. P-values determined by Kruskal-Wallis test. B) Disease progress curve of plants petiole-inoculated with UW551 cells stored in 20°C water microcosms (light green) vs. traditional frozen stocks (dark green). P-value determined by two-way repeated measures ANOVA. C) Disease progress curve of plants petiole-inoculated with UW770 cells stored in 20°C water microcosms (light orange) vs. traditional frozen stocks (dark orange). P-value determined by two-way repeated measures ANOVA.

### Tomato plants inoculated with UW770 are commonly asymptomatic carriers

To test virulence, we similarly inoculated plants via the petiole and quantified virulence for 14 days. Rapid disease progression and complete plant death was observed for UW551-infected plants (Figure 4B). However, there was a slight but significant delay in disease progression for water-adapted UW551 as compared to cells from typical laboratory storage conditions. Modest disease was observed for plants inoculated with UW770 (Figure 4C). Disease progression of plants infected with water-adapted UW770 cells significantly differed from laboratory-stored cells: there was a delay in initial onset, but a higher disease burden during the last few days of plants infected with water-adapted cells. Intriguingly, despite the modest symptoms, we noted that nearly all inoculated tomato plants were successfully colonized at 14 days post-infection. At the conclusion of the 14-day disease monitoring window, each tomato plant was de-topped above the site of inoculation and “stamped” onto an agar plate to qualitatively evaluate colonization. We found that while only 38% of plants inoculated with the laboratory-stored UW770 showed visible symptoms, 95% had detectable stem populations after 14 days. Similarly, while 53% of plants inoculated with the water-adapted UW770 showed visible symptoms, 90% had detectable stem populations after 14 days. We also concentrated water samples from the UW551 and UW770 12°C and 4°C microcosms and inoculated susceptible tomato plants. No tomato plants inoculated with these samples exhibited any symptoms, nor had detectable stem populations after 14 days (data not shown).

## Discussion

Plant-pathogenic *Ralstonia* concerningly persists in surface water (Vogelaar *et al*. 2023). Agricultural run-off can mobilize this pathogen from terrestrial to aquatic systems, and some hosts, such as the semi-aquatic weed *Solanum dulcamara*, are known to provide a reservoir for overwintering and proliferation within these environmental systems. Surface water contamination complicates management strategies and presents unique challenges for eradication efforts. It is therefore critical to understand *Ralstonia* dynamics in aquatic ecosystems to better inform management. We selected three temperatures that represent a range of environmentally-relevant conditions plant-pathogenic *Ralstonia* cells may experience in European surface waterways throughout the year (van Overbeek *et al*. 2004; Caruso *et al*. 2005).

Numerous studies have investigated how plant-pathogenic *Ralstonia* persists in aquatic microcosms. These studies support an important role for temperature in dictating behavior in water (van Elsas *et al*. 2001; Álvarez, López and Biosca 2022). Generally, *Ralstonia* maintains stable and culturable populations at warmer temperatures, while a viable but non-culturable (VBNC) state is induced at cold temperatures (4**°**C) (Milling *et al*. 2009). Our results are in line with these previous reports. Though we did not explicitly measure the viability of cells in the 4**°**C treatment, we observed a loss of culturability that is consistent with the VBNC state. We found that the IIB-1 strain UW551 remained culturable at cooler temperatures for longer than UW770. This strain may therefore be more resistant to cold temperatures in aquatic environments, and thus persist in waterways throughout seasonal fluctuations.

Multiple studies have shown that *Ralstonia* persists longer in sterile water than non-sterile water, likely due to negative interactions with native microbiota (van Elsas *et al*. 2001; Alvarez, López and Biosca 2007). While we used sterilized river water for our microcosms, we inoculated strains singly or dually to assess the impact of species interactions on population dynamics. We observed no major difference in each strain’s fitness between mono- and dual-inoculated treatments. This suggests that if the strains are interacting in this environment, the outcome does not majorly affect culturable cell abundance over time. However, we cannot rule out other antagonistic or beneficial interactions under different environmental conditions. In fact, our *in planta* data suggests the strains interact differently within a host.

We compared the host interaction phenotypes of *Rs* and *Rps* cells incubated at 20**°**C in water for 25 weeks to cells stored in typical lab conditions (frozen glycerol stocks and short-term growth in rich media). While water-adaptation modestly altered host colonization and virulence patterns for each strain, the biological outcome was generally the same: cells colonized stems and caused (a)symptomatic disease. This suggests water-adaptation does not significantly change plant interaction outcomes - at least for aquatic microcosm cells that were still culturable. Our non-culturable water microcosm treatments (4**°**C and 12**°**C) did not cause any observable disease or measurably colonize stems. In some instances, VBNC cells from 4**°**C water microcosms have been shown to cause disease when introduced directly into plant tissue (Álvarez, López and Biosca 2022). However, we used cells that had been incubated at these temperatures for much longer (25 weeks vs. 5-6 weeks) and only measured disease/colonization after 2 weeks, rather than 4 weeks.

Plant inoculations using traditionally prepared *Ralstonia* cultures revealed that in mono-inoculation, UW770 colonized tomato stems at higher levels than UW551. However, this is in contrast with the virulence curves generated from similar inoculations: 100% of plants succumbed to UW551 infection within 9 days, while only ∼45% of plants even showed symptoms when infected with UW770. Interestingly, despite the moderate symptom development, nearly all of the UW770-infected tomato plants had detectable populations after 2 weeks. This large number of asymptomatic carriers is important to consider for management practices, and is in line with the repeated introduction of phylotype I strains into European markets in latently infected plants.

During competitive co-inoculations, UW770 consistently colonized stem tissue at a much higher level than UW551. The initial experimental design inoculated approximately 1,000 cells of each strain into stem tissue simultaneously. When we altered the inoculum to give UW551 a larger proportion of the initial cells, we observed a higher colonization level of UW551 three days later. However, this could be due to the stochasticity of such a small number of UW770 cells (∼20) being inoculated. Therefore, if these two strains are given equivalent opportunities to colonize the same plant in warm temperatures, UW770 may be the dominant pathogen. These co-inoculations were performed at 28**°**C. Since UW551 exhibits cool virulence (Dewberry *et al*. 2024), future studies investigating host interactions at cooler temperatures may reveal interesting temperature-dependent competitive outcomes in plants.

Further, our data supports that the environment dictates the interactions between these two plant-pathogenic *Ralstonia* strains. UW770 negatively affected the cell count of UW551 in plant tissue, but not water microcosms. This could be due to indirect or direct competition between the strains. For example, UW770 grows to a higher level in plant stems after 3 days in mono-inoculation. It may therefore have an increased ability to uptake nutrients and grow to high cell densities in the xylem as compared to UW551. In co-inoculations, UW770 may therefore prevent UW551 from growing by more quickly using available nutrients. However, these results could also be explained by more direct antagonistic interactions, such as through the production of bacteriocins. Previous work has shown that some members of the RSSC produce bacteriocins that target other plant-pathogenic *Ralstonia (Huerta, Milling and Allen 2015)*. While we did not observe any evidence for direct bacterial antagonism *in vitro*, we can not rule this out, as bacteriocin production can be context-dependent. In fact, our different interactions in aquatic systems and in the plant suggest this is the case.

In this work, we investigated the ecological interactions of two quarantine-concern plant-pathogenic *Ralstonia* strains in river water and tomato plants. We uniquely look at the interaction between two naturally co-existing *Ralstonia* strains. It is critical to consider the intersection of temperature and virulence of plant-pathogenic *Ralstonia* as the climate continues to warm. Together, our findings show that interactions between *Rs* and *Rps* strains are context-dependent, strains maintain virulence after 6 months in 20**°**C water, and I-33 strains may outcompete IIB-1 strains in susceptible host environments.

## Acknowledgements

We kindly thank Jan van der Wolf and Maria Bergsma-Vlami (Wageningen University) for generously sharing the rifampin-resistant I-33 strain isolated from the Netherlands.

## Supplemental Figures/Tables

**Figure S1.**
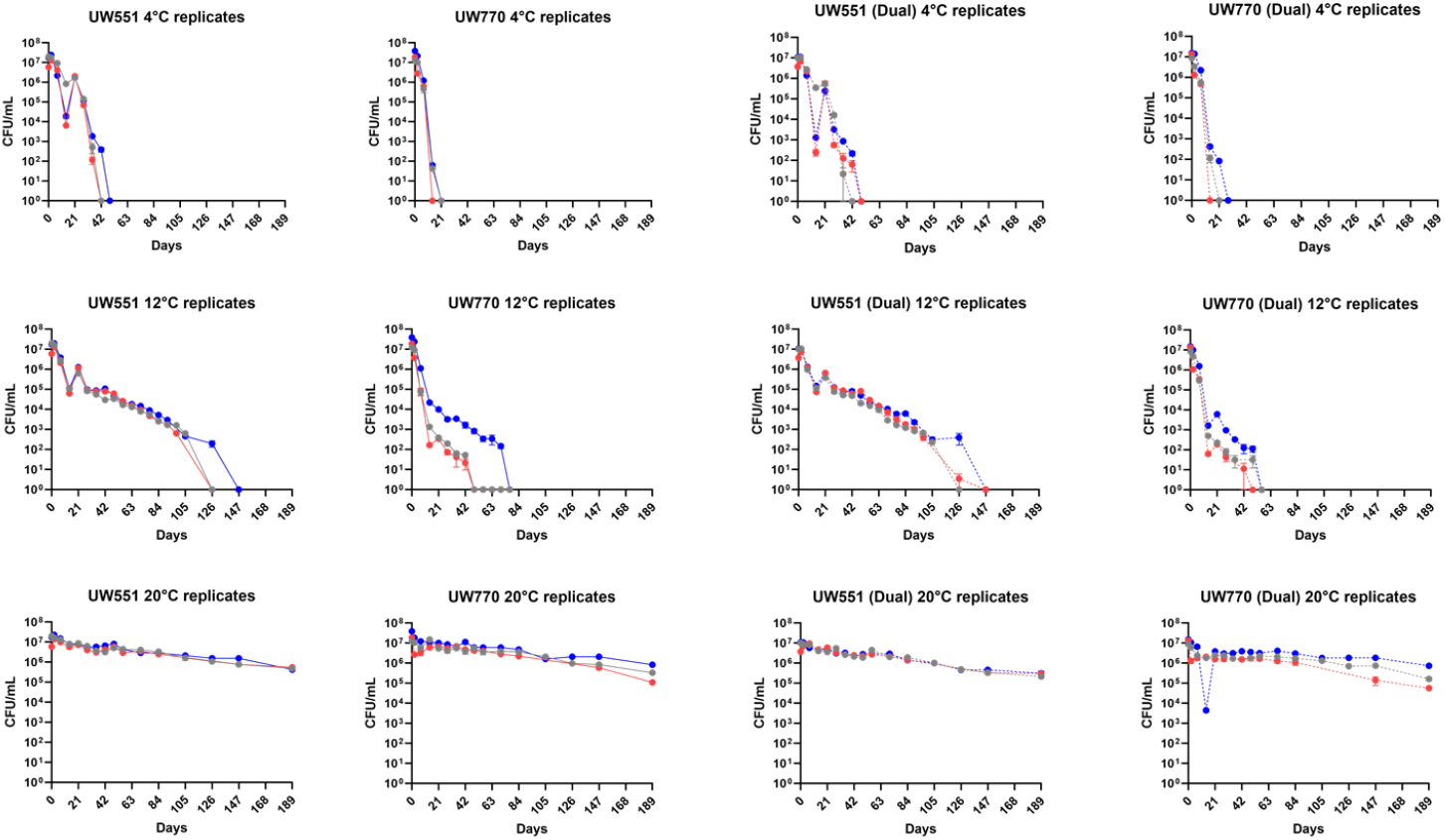
*Ralstonia* culturability across individual replicates within aquatic microcosm treatments. UW551 and UW770 were inoculated singly (mono) or dually (dual) into river water microcosms and incubated at either 4°C, 12°C, or 20°C for 6 months. Lines are colored based on replicate: gray lines = replicate 1, red lines = replicate 2, blue lines = replicate 3. Lines represent the average of 3 individual tubes for 3 biological replicates (n=9). Bars represent the SEM at each timepoint. The “Dual” lines represent the CFUs of each respective strain from the dual-inoculated microcosms. A single sub-sample from the dual-inoculated tubes was dilution plated on CPG + appropriate antibiotic to enumerate each strain individually. Plates used for sampling after 21 days also contained catalase. The panels represent an increasing temperature gradient from left to right.

**Table S1.** Concentration of cations, anions, and organic/inorganic carbon in water samples.

|  |  | Sample |  |  |
| --- | --- | --- | --- | --- |
|  |  | Rep 1 | Rep 2 | Rep 3 |
| <u>Anions (mg/L)</u> | Fluoride | 0.06 | 0.06 | 0.06 |
|  | Chloride | 55.39 | 55.66 | 56.18 |
|  | Nitrite | 0.05 | 0.22 | 0.11 |
|  | Bromide | 0.03 | 0.03 | 0.03 |
|  | Nitrate | 3.07 | 5.54 | 4.62 |
|  | Phosphate | n.a. | n.a. | n.a. |
|  | Sulfate | 17.37 | 17.56 | 19.10 |
| <u>Cations (mg/L)</u> | Potassium | 1.88 | 2.00 | 2.08 |
|  | Sodium | 24.05 | 24.17 | 24.07 |
|  | Ammonium | 0.19 | 0.31 | 0.22 |
|  | Magnesium | 37.67 | 38.60 | 40.58 |
|  | Calcium | 14.92 | 16.34 | 10.12 |
| <u>Carbon (mg/L)</u> | Organic C | 5.59 | 5.47 | 4.60 |
|  | Inorganic C | 34.36 | 33.90 | 32.50 |

**Figure S2.**
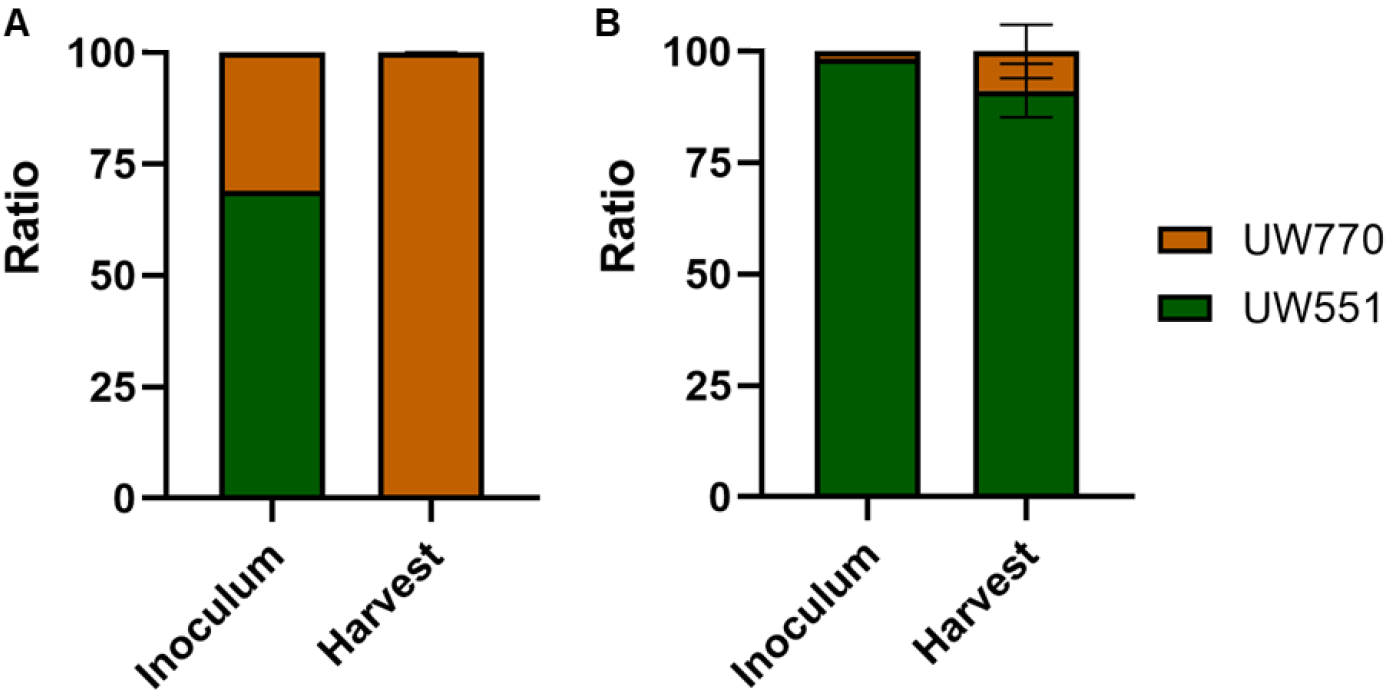
Stem colonization ratio of UW551 and UW770 following varied inoculum ratios. The inoculum ratio was adjusted to increase the number of UW551 cells compared to UW770 cells. A) Plants inoculated with ∼70% UW551 cells were still dominated by UW770 at 3 days post-inoculation. B) Plants inoculated with ∼98% UW551 cells were dominated by UW551 at 3 days post-inoculation. Data in each panel represents a single replicate with 12 plants.

